# Expression of XCR1, CX3CR1, CSF1R and ADGRE1 on subsets of bovine blood dendritic cells and monocytes

**DOI:** 10.1101/2024.09.17.613502

**Authors:** Stephanie C Talker, Zhiguang Wu, Inga Dry, Artur Summerfield, Jayne C Hope

**Affiliations:** Institute of Virology and Immunology, Bern, Switzerland; Department of Infectious Diseases and Pathobiology, Vetsuisse Faculty, University of Bern, Bern, Switzerland; The Roslin Institute, University of Edinburgh, Easter Bush, EH25 9RG, UK

**Author notes:** **Correspondence:** Stephanie C. Talker, Institute of Virology and Immunology, Bern Switzerland. Shared senior authorship (Artur Summerfield, Jayne C Hope).

**Keywords:** dendritic cells, monocytes, cattle, phenotyping, flow cytometry

## Abstract

Among dendritic cells (DC) and monocytes in blood of cattle we can distinguish conventional and plasmacytoid DC (cDC1, cDC2, pDC) and classical, intermediate and nonclassical monocytes (cM, intM, ncM), respectively. Phenotypic definitions of subsets rely on combinations of only a few markers such as CD13 and CD4 for Flt3^+^ dendritic cells, and CD14 and CD16 for CD172a^high^ monocytes. Additional reagents for flow cytometry are urgently needed to refine these phenotypic classifications and account for heterogeneity of these cells, in particular cDC2 and intM.

In the current study we have investigated expression of CX3CR1 and ADGRE1 on bovine blood DC and monocyte subsets, and have tested two newly generated bovine recombinant proteins (XCL1 and CSF1) for staining of XCR1 and CSF1R.

Staining patterns obtained by multicolor flow cytometry of peripheral blood mononuclear cells from two cows were largely in line with gene expression data available from a previous study (bulk RNA-seq of sorted DC and monocyte subsets from bovine blood).

## Introduction

Dendritic-cell and monocyte subsets in blood of cattle can be defined by relatively simple flow cytometry panels that have been confirmed by bulk RNA-seq of sorted subsets (Talker et al., 2018). Within Flt3^+^ cells, cDC1 can be identified as CD13^+^, pDC as CD4^+^, and cDC2 as negative for both markers. Especially cDC2 are expected to be far more heterogeneous, which is also reflected in the heterogeneous expression of most molecules analyzed in phenotyping experiments (Talker et al., 2018). Similarly, intermediate monocytes, gated as CD14^+^CD16^+/high^ within CD172a^high^ cells, likely contain mixed populations, potentially with opposing functions (Talker et al., 2022).

Thanks to joint efforts of the veterinary immunology community, the immunological toolbox for cattle is continuously increasing (Entrican et al., 2020; Mwangi et al., 2020), enabling extensive phenotyping of immune-cell subsets (www.immunologicaltoolbox.co.uk).

Recently, two additional reagents were described enabling the staining of CX3CR1 and ADGRE1 on bovine cells (Randall et al., 2024). Here, we employ these reagents to phenotype above-mentioned DC and monocyte subsets in bovine blood.

Additional reagents generated at the Roslin institute and tested in the present study target bovine XCR1 and CSF1R, two key markers for cDC1 and monocytic cells, respectively.

With bulk RNA-seq data available for all above-mentioned DC and monocyte subsets (Talker et al., 2018), we compared staining patterns obtained by flow cytometry to molecule-specific transcript levels in respective subsets.

As reported here, flow cytometry analysis for two cows supports specificity of all reagents, although unspecific staining needs to be taken into consideration, in particular when dealing with a high number of acquired events for analysis of DC subsets.

## Materials and Methods

### Generation of bovine XCL1 and CSF1

The cDNA coding for bioactive region (mature protein 3-225) of bovine CSF1 (NCBI Reference Sequence: NM_174026.1) with sequences for a 6xHIS and a FLAG at N terminal of CSF1 was synthesised by INTEGRATED DNA TECHNOLOGIES (IDT) and subsequently subcloned in frame into pFUSEN-hG1Fc (InvivoGen) using BsiWI and NheI restriction enzyme sites. This recombinant CSF1 has a Human IgG1-Fc tag (hG1Fc) at N-terminal, followed by a 6xHIS-linker and a FLAG tag in between to allow cleavage of hG1Fc when required.

The cDNA for bovine XCL1 (NCBI Reference Sequence: NM_175716) was synthesised by IDT and subsequently subcloned in frame into pKW06 vector (Wu et al., 2016) to express recombinant protein with a C-terminal human IgG1 Fc tag (XCL1-Fc), using NheI and BglII restriction enzyme sites.

Both recombinant XCL1 and CSF1 were produced by expi293TMexpression system (ThermoFisher SCIENTIFIC) and purified using a HiTrap Protein G HP column (Cytiva) as per instruction. Protein identity and purity was proved by mass spectrometry (MS), carried out by the Proteomics and Metabolomics Facility at the Roslin Institute, University of Edinburgh.

### Isolation of peripheral blood mononuclear cells (PBMC)

Blood collection and PBMC isolation were performed as previously described (Talker et al., 2018). The blood sampling was performed in compliance with the Swiss animal protection law and approved by the animal welfare committee of the Canton of Bern, Switzerland, license number BE42/2021.

### Flow cytometry

Staining of freshly isolated PBMC was performed in 96-well plates (U-bottom) with 1-2 × 10^7^ cells per sample. All steps were performed on ice or at 4 °C. The procedure encompassed five incubation steps à 15 minutes, including a pre-incubation with purified bovine IgG (Bethyl laboratories, Montgomery, USA) to block Fc receptors, and a blocking step with mouse IgG (Jackson ImmunoResearch, Switzerland) in order to block remaining binding sites of secondary antibodies. Washing steps between incubations were done with Cell Wash (BD Biosciences, Allschwil, Switzerland). Staining panels are given in Table 1. Bovine Flt3L-His (NCBI NM_181030.2) was produced as previously described (Ziegler et al., 2016).

**Table 1.**
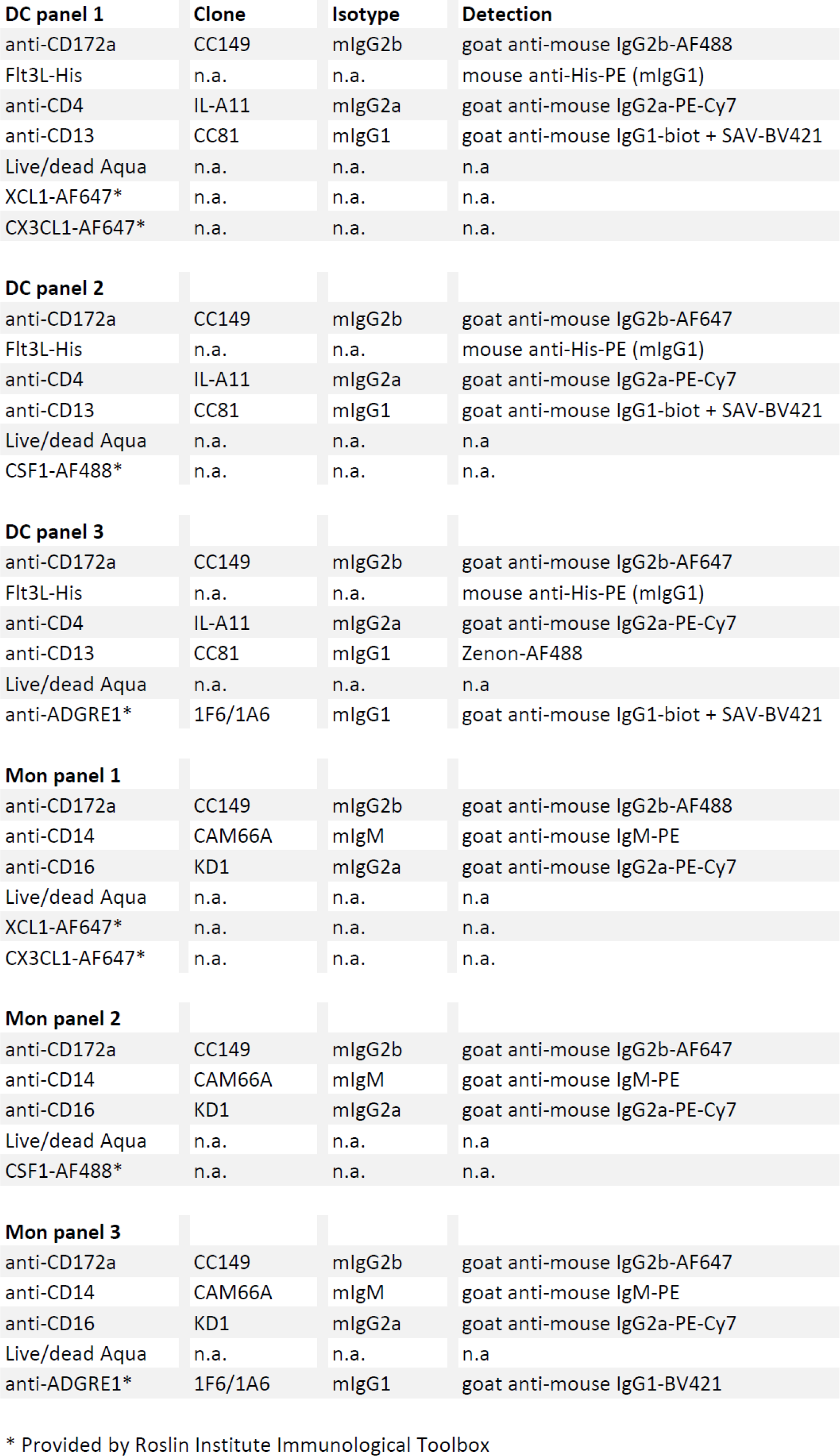
Staining panels.

**Staining panels**
| **DC panel 1** | **Clone** | **Isotype** | **Detection** |
| --- | --- | --- | --- |
| anti-CD172a | CC149 | mIgG2b | goat anti-mouse IgG2b-AF488 |
| Flt3L-His | n.a. | n.a. | mouse anti-His-PE (mIgG1) |
| anti-CD4 | IL-A11 | mIgG2a | goat anti-mouse IgG2a-PE-Cy7 |
| anti-CD13 | CC81 | mIgG1 | goat anti-mouse IgG1-biot + SAV-BV421 |
| Live/dead Aqua | n.a. | n.a. | n.a. |
| XCL1-AF647* | n.a. | n.a. | n.a. |
| CX3CL1-AF647* | n.a. | n.a. | n.a. |
| **DC panel 2** |  |  |  |
| anti-CD172a | CC149 | mIgG2b | goat anti-mouse IgG2b-AF647 |
| Flt3L-His | n.a. | n.a. | mouse anti-His-PE (mIgG1) |
| anti-CD4 | IL-A11 | mIgG2a | goat anti-mouse IgG2a-PE-Cy7 |
| anti-CD13 | CC81 | mIgG1 | goat anti-mouse IgG1-biot + SAV-BV421 |
| Live/dead Aqua | n.a. | n.a. | n.a. |
| CSF1-AF488* | n.a. | n.a. | n.a. |
| **DC panel 3** |  |  |  |
| anti-CD172a | CC149 | mIgG2b | goat anti-mouse IgG2b-AF647 |
| Flt3L-His | n.a. | n.a. | mouse anti-His-PE (mIgG1) |
| anti-CD4 | IL-A11 | mIgG2a | goat anti-mouse IgG2a-PE-Cy7 |
| anti-CD13 | CC81 | mIgG1 | Zenon-AF488 |
| Live/dead Aqua | n.a. | n.a. | n.a. |
| anti-ADGRE1* | 1F6/1A6 | mIgG1 | goat anti-mouse IgG1-biot + SAV-BV421 |
| **Mon panel 1** |  |  |  |
| anti-CD172a | CC149 | mIgG2b | goat anti-mouse IgG2b-AF488 |
| anti-CD14 | CAM66A | mIgM | goat anti-mouse IgM-PE |
| anti-CD16 | KD1 | mIgG2a | goat anti-mouse IgG2a-PE-Cy7 |
| Live/dead Aqua | n.a. | n.a. | n.a. |
| XCL1-AF647* | n.a. | n.a. | n.a. |
| CX3CL1-AF647* | n.a. | n.a. | n.a. |
| **Mon panel 2** |  |  |  |
| anti-CD172a | CC149 | mIgG2b | goat anti-mouse IgG2b-AF647 |
| anti-CD14 | CAM66A | mIgM | goat anti-mouse IgM-PE |
| anti-CD16 | KD1 | mIgG2a | goat anti-mouse IgG2a-PE-Cy7 |
| Live/dead Aqua | n.a. | n.a. | n.a. |
| CSF1-AF488* | n.a. | n.a. | n.a. |
| **Mon panel 3** |  |  |  |
| anti-CD172a | CC149 | mIgG2b | goat anti-mouse IgG2b-AF647 |
| anti-CD14 | CAM66A | mIgM | goat anti-mouse IgM-PE |
| anti-CD16 | KD1 | mIgG2a | goat anti-mouse IgG2a-PE-Cy7 |
| Live/dead Aqua | n.a. | n.a. | n.a. |
| anti-ADGRE1* | 1F6/1A6 | mIgG1 | goat anti-mouse IgG1-BV421 |

Also, generation of bovine CX3CL1 and monoclonal antibody to ADGRE1 (clone number 1F6/1A6) were described previously (Randall et al., 2024). Recombinant proteins XCL1, CX3CL1 and CSF1 were fluorescently labelled with Alexa Fluor (AF) 647 or AF488 using Alexa Fluor™ Antibody Labeling Kits (ThermoFisher SCIENTIFIC).

For every investigated marker (XCR1, CX3CR1, CSF1R, and ADGRE1) fluorescence-minus-one (FMO) controls were included to account for differences in autofluorescence between DC and monocyte subsets. Samples were acquired on a Cytek Aurora equipped with four lasers (V-B-YG-R). At least 1 × 10^6^ cells were acquired in the large-cell gate. Raw data was unmixed using single-stained samples as reference controls. Autofluorescence extraction was not performed. Large cells were gated in Spectroflo version 3.3.0 (Cytek Biosciences) and exported for further analysis in FlowJo version 10.10.0 (BD Biosciences).

## Results & Discussion

To investigate expression of XCR1, CX3CR1, CSF1R and ADGRE1 on bovine DC and monocyte subsets, freshly isolated peripheral blood mononuclear cells (PBMC) were stained for multicolor flow cytometry. As evident from ungated plots shown in Figure 1, positive staining with bovine recombinant XCL1, CX3CL1, CSF1 and anti-ADGRE1 was not restricted to dendritic cells and monocytes. In particular bovine recombinant XCL1 stained a large proportion of Flt3^-^ cells. When markers were investigated on DC and monocyte subsets of interest, as illustrated in Figures 2 and 3, subset-specific staining patterns were observed. As expected, XCL1 stained bovine cDC1, indicative of specific binding to XCR1, while cDC2 remained unstained. However also pDC were partly stained by XCL1, which is surprising considering the lack of XCR1 transcripts in bovine pDC (Figure 4). Weak staining was also observed on cM, while intM and ncM were considered negative based on overlays with their corresponding FMO controls (Figure 3). Additional experiments are required to elucidate binding specificity of the bovine XCL1 protein described here. Certainly, judging from the experiments performed on PBMC of two cows, staining with XCL1 alone does not allow for gating on cDC1. Rather, this reagent may be used in combination with Flt3 or possibly CD13 or CD26. Moreover, based on recent findings in humans, XCR1 may only be expressed on a subset of cDC1, which appears to be more mature than cDC1 lacking XCR1 expression (Heger et al., 2023). Interesting in this regard, is also the observation that XCR1 expression was downregulated in bovine cDC1 stimulated with TLR ligands *in vitro* (Barut et al., 2020), suggesting that low XCR1 expression may be a feature of activated rather than immature cDC1, at least in cattle.

**Figure 1.**
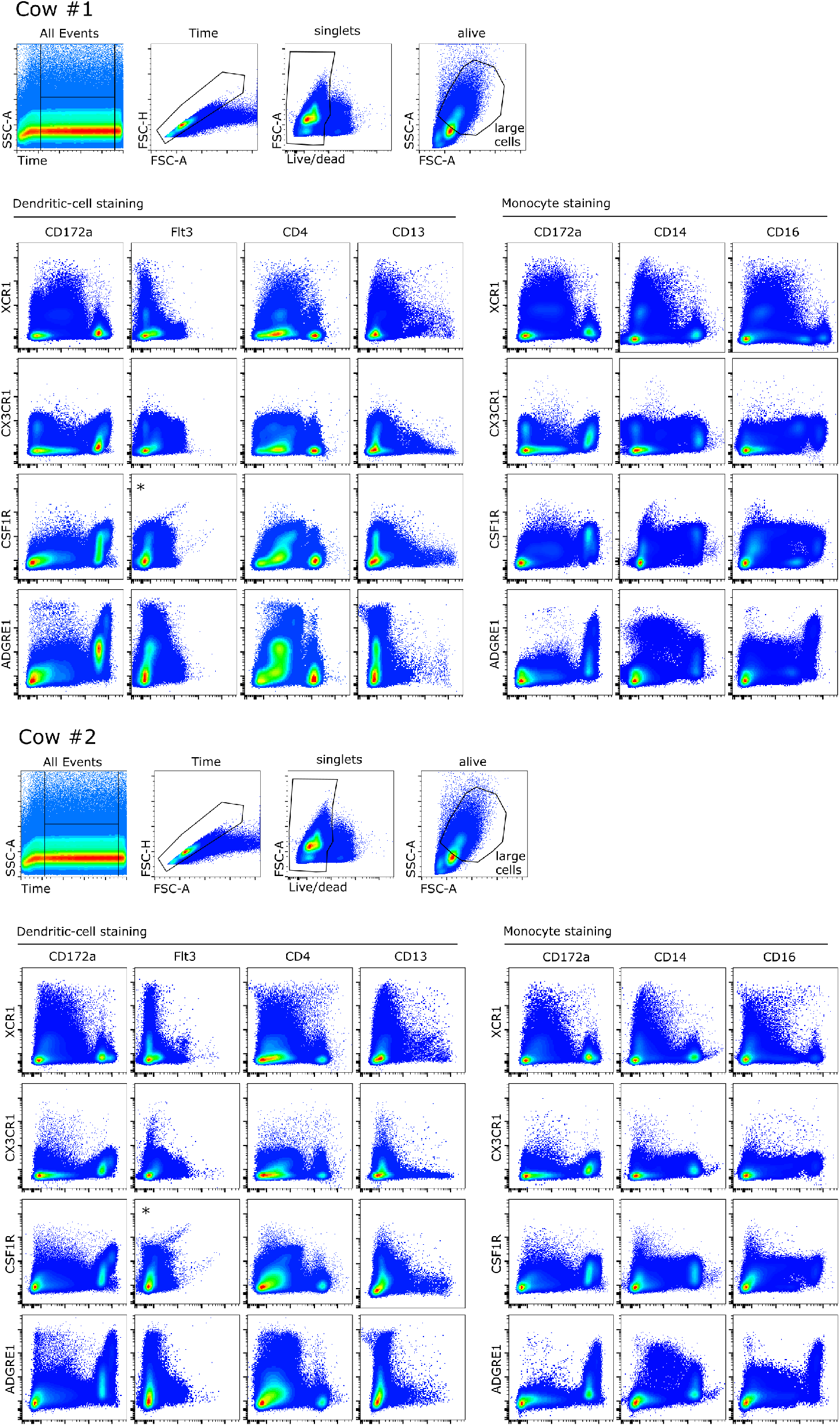
Staining patterns observed on ungated cells (large cells). Freshly isolated bovine PBMC were stained for expression of XCR1, CX3CR1, CSF1R and ADGRE1. Large cells were gated as indicated. Data for cow #1 (upper panel) and cow #2 (lower panel) is shown. Asterisks indicate plots that need to be interpreted with caution (anti-His PE theoretically binding to both Flt3L and CSF1).

**Figure 2.**
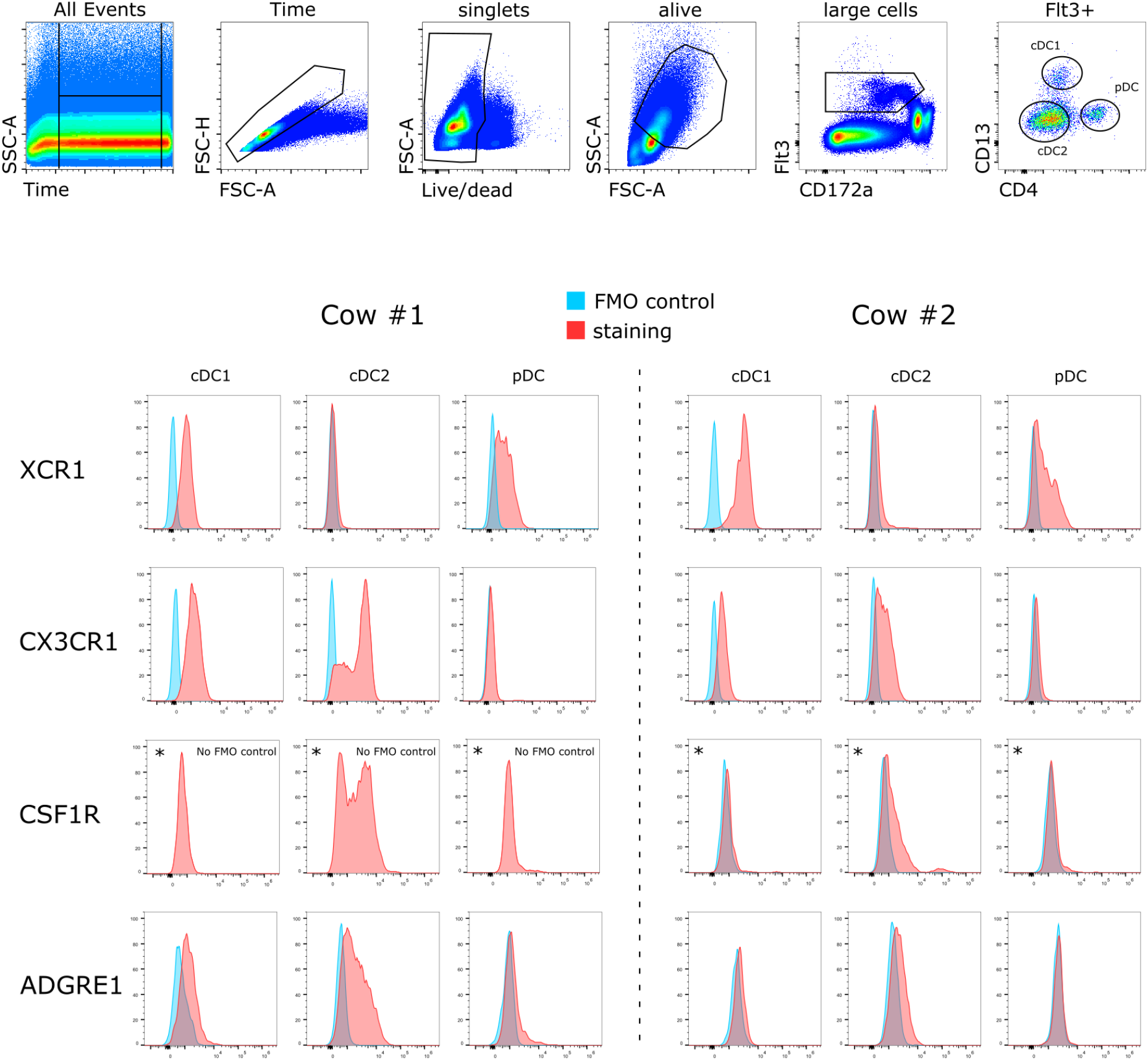
Phenotyping of DC subsets. Freshly isolated bovine PBMC (n=2) were stained for expression of XCR1, CX3CR1, CSF1R and ADGRE1. Dendritic-cell subsets were gated as Flt3^+^CD4^-^CD13^+^ (cDC1), Flt3^+^CD4^-^CD13^-^ (cDC2), and Flt3^+^CD4^+^CD13^-^ (pDC) from large cells (FSC-A vs. SSC-A) following exclusion of irregular events (Time), multiplets (FSC-A vs. FSC-H), and dead cells. For each cell subset, plots show overlays of FMO controls (blue histograms) and stained samples (red histograms). Asterisks indicate plots that need to be interpreted with caution (anti-His PE theoretically binding to both Flt3L and CSF1). Results for Flt3-independent gating of cDC2 are shown in Figure 3.

**Figure 3.**
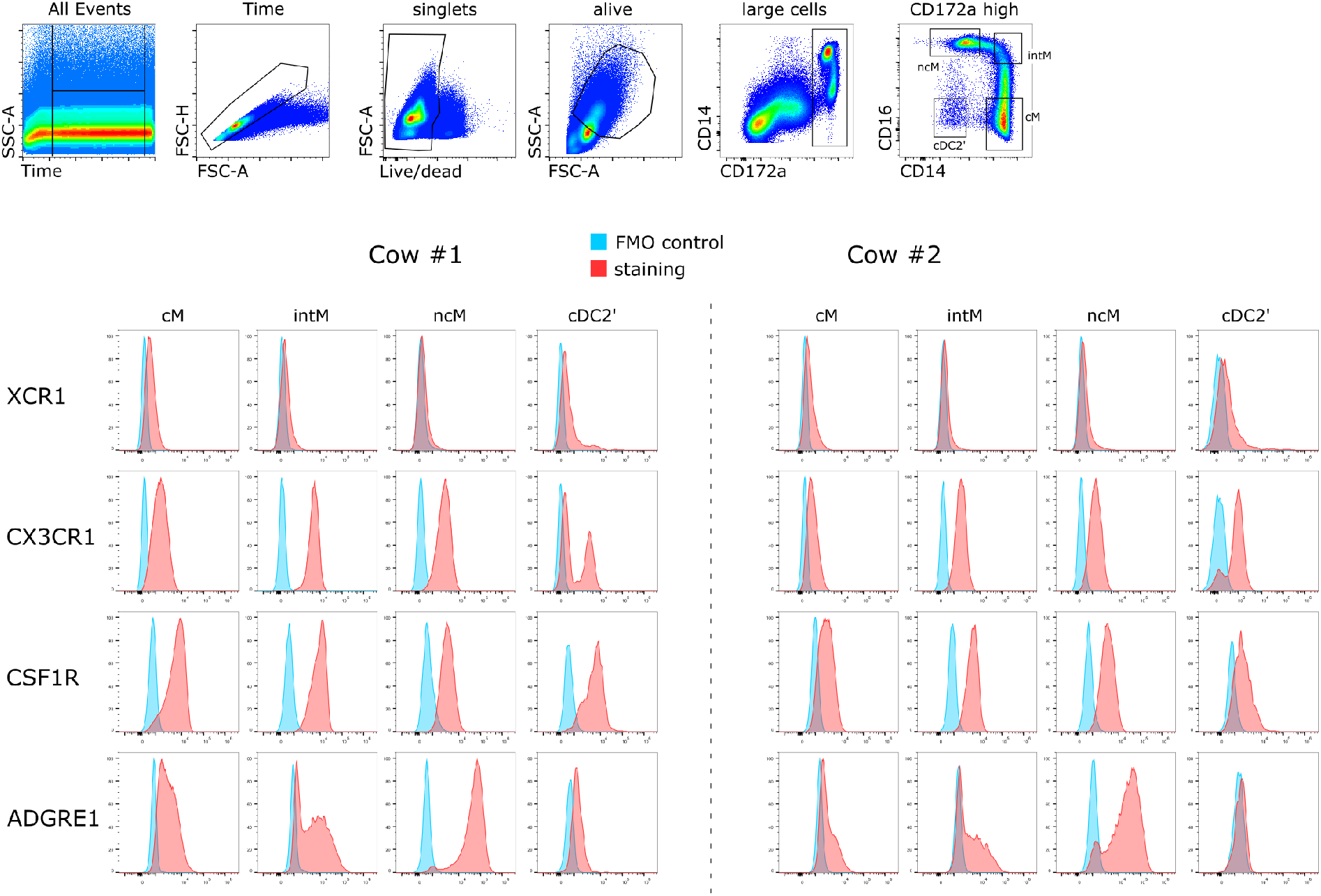
Phenotyping of monocyte subsets and cDC2’. Freshly isolated bovine PBMC (n=2) were stained for expression of XCR1, CX3CR1, CSF1R and ADGRE1. Monocyte subsets were gated as CD172a^high^CD14^+^CD16^-^ (cM), CD172a^high^CD14^+^CD16^high^ (intM), and CD172a^high^CD14^-/dim^CD16^high^ (ncM), and cDC2’ were gated as CD172a^high^CD14^-^CD16^-^ within large cells (FSC-A vs. SSC-A) following exclusion of irregular events (Time), multiplets (FSC-A vs. FSC-H), and dead cells. For each cell subset, plots show overlays of FMO controls (blue histograms) and stained samples (red histograms).

**Figure 4.**
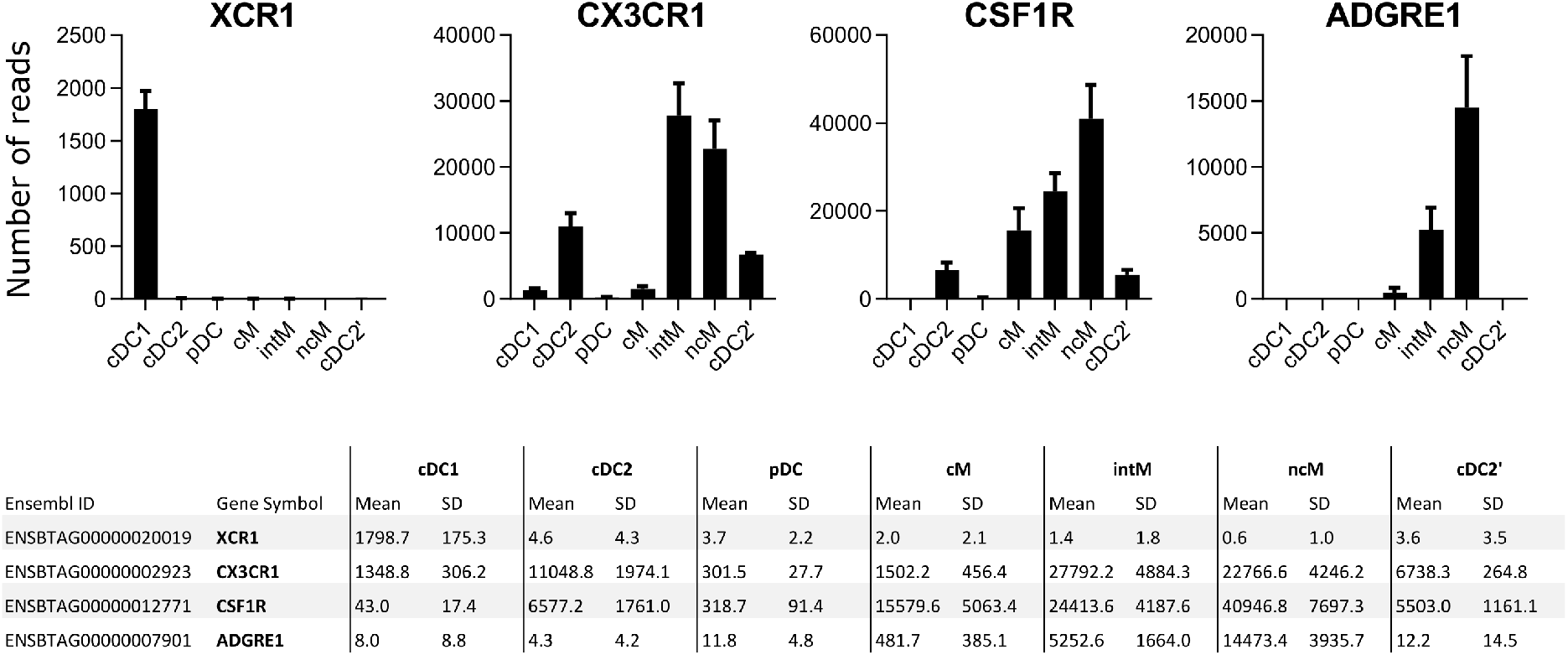
Expression levels of XCR1, CX3CR1, CSF1R and ADGRE1 as determined by bulk RNA-seq of DC and monocyte subsets sorted from bovine blood (Talker et al., 2018). Number of reads are shown as mean and standard deviation (n=3).

Transcription of CX3CR1 is reported to allow for distinction of ncM from cM in both human and cattle (Corripio-Miyar et al., 2015; Talker et al., 2022). Accordingly, ncM displayed higher levels of CX3CL1 staining than cM, even if transcriptomic data (Figure 4) would have suggested a more pronounced difference between these subsets. Both transcriptomics and CX3CL1 staining suggest that intM express the highest level of CX3CR1. Among DC, both cDC1 and cDC2 were stained with CX3CL1, while pDC were not, which is in line with transcriptomic data. Notably, one of the investigated animals contained two subsets of cDC2 with differing CX3CR1 expression levels. The CX3CR1 expressing subset may potentially be related to the CX3CR1^+^ cDC2 subset described for mice (Minutti et al., 2024).

The receptor for CSF1 (CSF-1R) is described as a marker for monocytic cells, with low levels of CSF1R transcripts detected also in bovine cDC2 (Figure 4). Accordingly, cDC1 and pDC remained unstained, and highest staining intensities were found for monocytes. Transcriptomics suggested increasing levels of CSF1R from cM via intM towards ncM, however this could not be confirmed by flow cytometry. Judging from the two investigated cows, expression levels on cM seem to vary depending on the animal, with a tendency towards highest expression on intM. Notably, two subsets of cDC2 with differing levels of CSF1R expression were apparent in cow#1 (compare with CX3CR1 expression above). It must be noted, however, that the 6xHis linker contained in recombinant CSF1 likely interfered with Flt3 staining (Flt3L-His + anti-His-PE). In fact, anti-His-PE and CSF1-AF488 were added to cells with the same mastermix, likely explaining reduced Flt3 staining in this sample (DC panel 2 shown in Table 1). Judging from the plots in Figure 1 however, erroneous co-labeling of Flt3^+^ cells with CSF1-AF488, as well as erroneous co-labeling of CSF1R^+^ cells with anti-His-PE appears to be negligible, potentially because AF488 molecules attached to CSF1 prevented binding of anti-His-PE by steric hindrance. Nevertheless, results obtained from DC panel 2 need to be interpreted with caution. Alternative gating of cDC2 as CD172a^high^CD14^-^CD16^-^ cells (cDC2’ in Figure 3) supports CSF1R expression on bovine cDC2. More stainings need to be performed to confirm these findings. An alternative staining panel might also solve the issue of relatively weak CSF1R staining in the current monocyte panel.

Expression of ADGRE1, a marker for monocytic cells across species also known as F4/80 (Waddell et al., 2018), was found predominantly on ncM and at lower levels on cM. Notably, two populations were evident within intM and to a lesser extent also in ncM, suggesting that ADGRE1 may be a suitable marker to further explore monocyte heterogeneity and the proposed transition of bovine cM to ncM (Talker et al., 2022). Transcriptomic data suggest that ADGRE1 is not expressed on DC (mean number of reads below 20), however cDC2 of cow #1 stained clearly positive (compare CX3CR1 and CSF1R expression on cDC2 of cow #1). As evident from ungated plots shown in Figure 1, some degree of unspecific ADGRE1 signal was present in the DC panel, likely due to the multi-step staining with anti-mIgG1-biotin and Streptavidin-BV421 (Table 1).

Taken together, we provide the first phenotyping of bovine DC and monocyte subsets with novel reagents provided by the Roslin Immunological Toolbox: bovine recombinant XCL1, CX3CL1, CSF1, and anti-bovine ADGRE1. Overall, we obtained staining patterns that are in line with transcriptomic data. For an updated version of the preprint and publication of a manuscript we aim to provide data on additional animals and alternative staining panels to corroborate these findings.

